# A preprocessing toolbox for 2-photon subcellular calcium imaging

**DOI:** 10.1101/2024.10.04.616737

**Authors:** Anqi Jiang, Chong Zhao, Mark Sheffield

## Abstract

Recording the spiking activity from subcellular compartments of neurons such as axons and dendrites during behavior with 2-photon calcium imaging is increasingly common yet remains challenging due to low signal-to-noise, inaccurate region-of-interest (ROI) identification, movement artifacts, and difficulty in grouping ROIs from the same neuron. To address these issues, we present a computationally efficient pre-processing pipeline for subcellular signal detection, movement artifact identification, and ROI grouping. For subcellular signal detection, we capture the frequency profile of calcium transient dynamics by applying Fast Fourier Transform (FFT) on smoothed time-series calcium traces collected from axon ROIs. We then apply band-pass filtering methods (e.g. 0.05 to 0.12 Hz) to select ROIs that contain frequencies that match the power band of transients. To remove motion artifacts from z-plane movement, we apply Principal Component Analysis on all calcium traces and use a Bottom-Up Segmentation change-point detection model on the first principal component. After removing movement artifacts, we further identify calcium transients from noise by analyzing their prominence and duration. Finally, ROIs with high activity correlation are grouped using hierarchical or k-means clustering. Using axon ROIs in the CA1 region, we confirm that both clustering methods effectively determine the optimal number of clusters in pairwise correlation matrices, yielding similar groupings to “ground truth” data. Our approach provides a guideline for standardizing the extraction of physiological signals from subcellular compartments during behavior with 2-photon calcium imaging.

## INTRODUCTION

Two-photon calcium imaging enables the recording of activity in subcellular neuronal structures such as dendrites, spines, and axons during behavior. Recent studies have demonstrated the feasibility of using this approach to investigate how specific sets of inputs transmit information to particular regions and how this information is integrated at postsynaptic sites[1-15]. This methodology requires meticulous analysis of calcium signals from these small structures, which are highly susceptible to movement artifacts and signal noise. Additionally, the intricate morphology of axonal and dendritic structures complicates the identification of segments belonging to the same neuron within the imaging plane. Addressing these challenges is essential for accurately interpreting the data collected in these experiments, which are increasingly being adopted by many laboratories. Standardizing the extraction of subcellular calcium signals through a widely accessible processing pipeline would facilitate direct comparisons across different studies.

Suite2P is the most popular software used for the initial identification of regions of interest (ROI) within a field of view (FOV)[16]. Suite2P can also be used for the initial identification of ROIs from a FOV containing subcellular structures such as axons. However, axon and dendritic ROIs are relatively small and are therefore subject to more noise than somas from movement artifacts and fewer pixels to extract signals from. Additional steps are therefore required following initial ROI identification by Suite2P to ensure ROIs contain real physiological signals that can then be analyzed. Various methods have been employed for the selection of active axon ROIs, such as manual labeling by visual inspection[1] or using convolutional neural network classifiers[12]. Manual labeling by looking through time-series movies for fluorescent axon or dendritic signals, while straightforward, is impractical for FOVs with many ROIs and is prone to user bias. Neural network classifiers, though effective, require extensive ground truth data for training, which is labor-intensive and computationally demanding, as well as being limited to one particular dataset, thus not generalizable.

Dual-color imaging offers another approach to identifying active subcellular ROIs by using a static fluorophore alongside an activity-dependent one[3, 9]. For instance, imaging calcium dynamics with GCaMP in a green channel and using mCherry in a red channel can help distinguish real calcium signals from noise. However, differences in fluorophore behavior and excitation/emission spectra means that subtracting the static channel from the active channel may not remove all noise and can introduce new signal artifacts. Employing dual lasers to optimize excitation could help with this but further complicates the process and the addition of second laser beam increases the probability of photo-damage to the tissue being imaged [17]. This is not to say a static channel is not useful, as it can help with motion correction of time-series FOVs in the X and Y axis and can help identify periods when the FOV may shift in the Z-axis, but simultaneous 2-channel imaging needs careful implementation and consideration.

Once subcellular ROIs are selected, identifying and correcting z-plane movement artifacts is critical. Slight brain movements during behavior can cause significant changes in the imaged structures, leading to false-positive calcium signals and this issue is amplified when imaging subcellular structures which are much smaller than somas[18]. Methods such as measuring pixel correlations within an ROI over time have been used to detect these artifacts, but they are computationally intensive and heavily dependent on the initial ROI selection[13, 14].

Recording from small structures like thin axons also presents a low signal-to-noise ratio challenge due to the limited number of pixels available for averaging physiological signals amidst inherent noise in the two-photon system[19]. Identifying which subcellular ROIs belong to the same neuron can improve the signal-to-noise ratio by providing more pixels for averaging. Further, because single axons and dendritic branches can come in and out of the FOV, the same neural structure can be represented by disconnected ROIs that are far apart in the FOV. If ROIs belonging to the same neuron are not grouped, they will be overrepresented in the data and can confound results. However, resolving which ROIs are part of the same structure remains a significant issue. Grouping ROIs that belong to the same neuron involves methods like hierarchical clustering and pairwise correlation thresholding. However, the lack of ground truth data to validate these groupings leaves their accuracy uncertain.

We present a preprocessing pipeline for identifying subcellular ROIs with real physiological signals, removing motion artifacts, and grouping ROIs belonging to the same neuron. We apply Fast Fourier Transform (FFT) on smoothed calcium traces to identify the frequency band of transients, followed by band-pass filtering to select ROIs matching the power band of transients. Principal Component Analysis (PCA) and Bottom-Up Segmentation change point detection are then used to remove motion artifacts. Transient peak detection based on prominence and duration identifies real calcium transients. Finally, we employ hierarchical or k-means clustering to group correlated ROIs, and validate this approach with “ground truth” axon clusters. Our method provides a guideline for analyzing axonal and dendritic activity using two-photon calcium imaging, aiding in the standardization of physiological signal extraction from subcellular structures during behavior.

## METHODS

### Experimental model and subject details

All experimental and surgical procedures were in accordance with the University of Chicago Animal Care and Use Committee guidelines. We used 10–20 week old C57BL/6-Tg(Grik4-cre)G32-4Stl/J mice (23–33 g). Mice were individually housed in a reverse 12 hour light/dark cycle and behavioral experiments were conducted during the animal’s dark cycle.

### Injection protocol

Mice were anesthetized and I.P. injected with 0.5ml of saline and 0.5ml of meloxicam. A small craniotomy was made over the right hippocampus CA3 region (2.0mm lateral, 1.7mm caudal respectively of Bregma). An axon-targeted genetically-encoded calcium indicator, AAV9-axon-GCaMP6s-P2A-mRuby3 (pAAV-hSynapsin1-axon-GCaMP6s-P2A-mRuby3 was a gift from Lin Tian Addgene viral prep # 112005-AAV9; http://n2t.net/addgene:112005; RRID:Addgene_112005) was injected (∼50 nL at a depth of 1.9 mm below the surface of the dura) using a beveled glass micropipette. Right after the viral injection, the site was covered with dental cement (Metabond, Parkell Corporation) and a metal head plate (Atlas Tool and Die Works). After ∼4-8 weeks, water scheduling (0.8-1ml per day) began. One week after the start of water scheduling, mice went through a hippocampal cannula implantation surgery over the right CA1 region (1.7mm lateral, −2.3mm caudal of Bregma). During the surgery, a head plate and head ring were attached to the cannula window for head-fixation and to house the microscope objective and block out ambient light. Post-surgery, mice had ad lib water, until a week later when they returned to water scheduling. Expression of axon-GCaMP6s in CA3 axons reached a steady state ∼6-12 weeks after the virus was injected, as monitored through 2P imaging.

### Two-photon imaging

Imaging was done using a laser scanning two-photon microscope (Neurolabware). Using a 8 kHz resonant scanner, images were collected at a frame rate of 15.49 Hz with unidirectional scanning through a 16×/0.8 NA/3 mm WD water immersion objective (MRP07220, Nikon). axon-GCaMP6s was excited at 920 nm with a femtosecond-pulsed laser (Insight DS+ Dual, Spectra-Physics) and emitted fluorescence was collected using two GaAsP PMTs (H11706, Hamamatsu). The average power of the laser measured after the objective ranged between 60–100 mW, and was kept constant across days of imaging. Time-series images were collected through Scanbox (Neurolabware) and a PicoScope Oscilloscope was used to synchronize frame acquisition timing with behavior.

### Image processing and ROI identification

Time-series images were preprocessed using Suite2p[16]. Movement artifacts in the X and Y-axis were corrected using rigid and non-rigid transformations based on the red mRuby channel. Regions of interest (ROIs) were also defined in the green channel using Suite2p. All identified ROIs were fed into SUBPREP for subsequent ROI selection. Raw fluorescence traces from ROIs were first smoothed using a Savitzky-Golay filter.

## RESULTS

### Frequency-based approach for transient identification and subcellular ROI selection

In selecting subcellular ROIs that contain real calcium transients, we developed a frequency-based ROI selection pipeline. The key idea underlying our selection procedure was that the rise-to-peak and decay phase of real transients have unique frequency domains. In particular, the rise of the transient occurs within a faster frequency band than the decay of the transient. Therefore, we designed a filter that captures both the rise and the decay of transients. To determine the key frequencies for the rise and the decay, we calculated the frequency bands using example ROIs with clear visible transients and high signal to noise ratio (SNR) and compared to the frequency bands from all ROIs. Because the vast majority of ROIs from Suite2P (85-95%, depending on the dataset) contain only noise, example ROIs with clear transients returned higher power in the frequency bands that reflect the rise and decay of transients compared to the mean of all ROIs. The frequency band with elevated power from example ROIs with clear transients became our frequency band of interest. If the filtered signal from an ROI returned high power in our frequency band of interest, the ROI would be selected as “likely to contain transients”. By contrast, if the filtered signal did not contain sufficient power (see below) in our frequency of interest, the ROI would be marked as “not likely to contain transients”. Note that the specific frequency band containing real transients will depend on which calcium indicator is being imaged (e.g. GCaMP6s versus GCamP8f) and which subcellular structures are being imaged (e.g. axons versus dendrites). The key frequency band will therefore need to be determined for each user dependent on the specifics of the dataset.

To quantify the power within the key frequency band among all subcellular ROIs, we applied a band-pass filter using the key frequency range on smoothed time-series calcium traces using the filtfilt function in the python script package. We utilized a third-order Butterworth filter, a type of infinite impulse response filter known for its maximally flat frequency response within the passband. By selecting this design of our filter, we ensured effective attenuation of frequencies outside the passband while minimizing phase distortion within the passband. First, a low-pass filter with a cutoff frequency of 0.13 Hz (or the higher bound of key frequency bands from example ROIs) was applied to the green channel signal to remove any high-frequency noise. Then, a high-pass filter with a cutoff frequency of 0.03 Hz (or the lower bound of key frequency bands from example ROIs) was applied to remove any low-frequency noise (e.g. Z-plane drift can be a source of low frequency noise). In particular, the filtfilt function employed a zero-phase digital filtering technique, which applied the filter forward and then backward to eliminate phase distortion, resulting in a zero-phase delay. Our procedure ensured that the filtered signal retains its temporal integrity. By using filtfilt, the signal was effectively band-pass filtered, removing both high-frequency noise and low-frequency drift, while preserving the original signal’s phase and shape.

Once the signal had been band-pass filtered using the filtfilt function, the power within the frequency band of interest (e.g. 0.03 Hz to 0.13 Hz) could be calculated. Then, we calculated the normalized power of the signal such that the power of interest was divided by the total power of the signal. An advantage of using normalized power was that the range was bounded with an upper limit of 1 and a lower limit of 0. Therefore, normalized power enabled comparisons across different traces and different animals, while absolute power was unbounded and depends on the length of the signal. If the power within this frequency band exceeded a predefined threshold (e.g. 0.3 of the normalized power), the ROI was defined as containing transients and the ROI selected. Using a higher threshold (e.g. 0.5) returns fewer ROIs but with higher SNR. This approach allowed for the identification and isolation of signals that contain significant activity within the specified frequency range, facilitating the extraction of ROIs with transient-like patterns within their traces.

**Fig. 3** shows two example ROIs from our dataset. The green ROI in **Fig. 3A** had a relatively low power in the frequency band of interest shown in **Fig. 3B** (0.182), suggesting this ROI is unlikely to contain transients. Contrarily, the black axon in **Fig. 3A**, which shows clear and obvious transients, had a higher power in the frequency range of interest (**Fig. 3B**, black; 0.454). Applying this approach to all ROIs from an example mouse, the distribution of the normalized power is shown in **Fig. 3C**. The ROIs selected as “likely to contain transient” were the ones with power above 0.3 (red dotted line) on the right side of the histogram that were kept for the next steps of the analysis, and the ROIs marked as “not likely to have transient” were shown on the left of the dotted line that were not used in the remainder of the analysis pipeline.

**Fig. 1.**
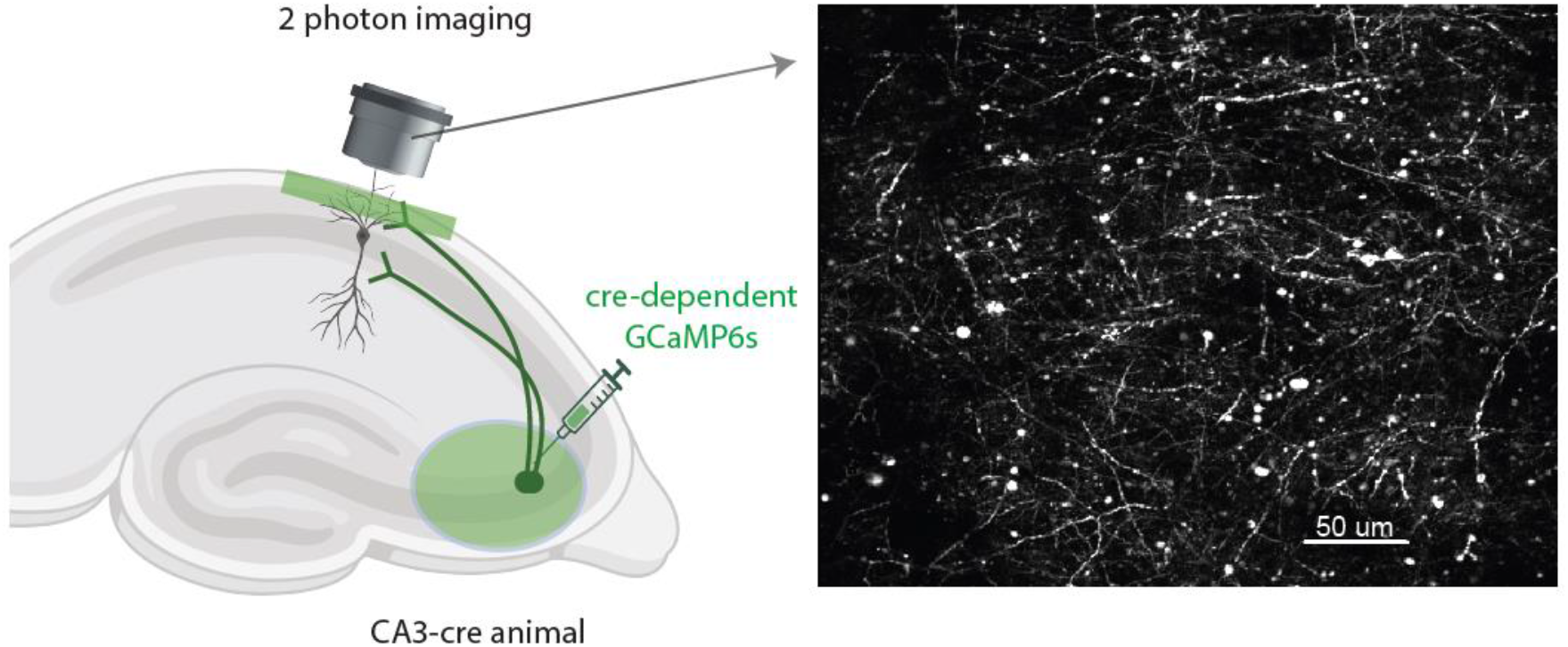
Imaging CA3 axons in CA1. Left: Schematic representation of CA3 axonal imaging in CA1 Stratum Oriens. Right: Example CA1 field of view containing CA3 axons. Image is the max projection from 5000 frames.

**Fig. 2:**
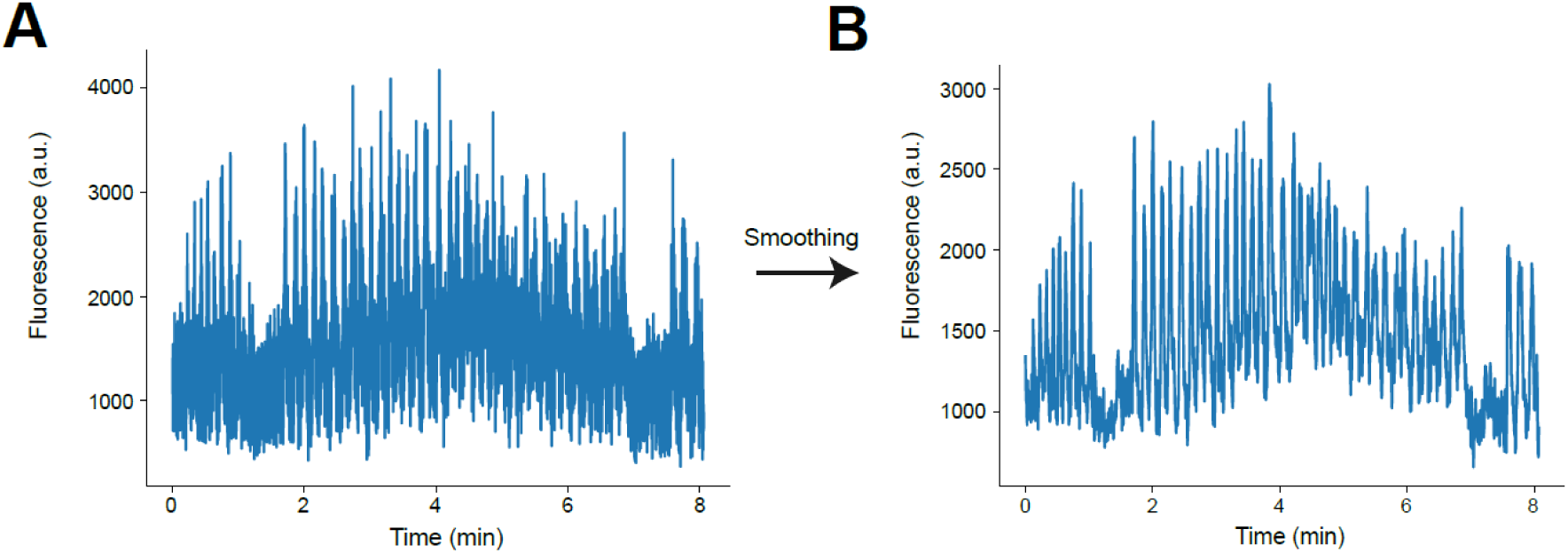
Example time-series fluorescent traces from CA3 axons in CA1 before and after smoothing. **A**. Example raw fluorescent time-series trace from an ROI identified by Suite2P. **B**. The same trace after smoothing using the Savitzky-Golay filter.

**Fig. 3.**
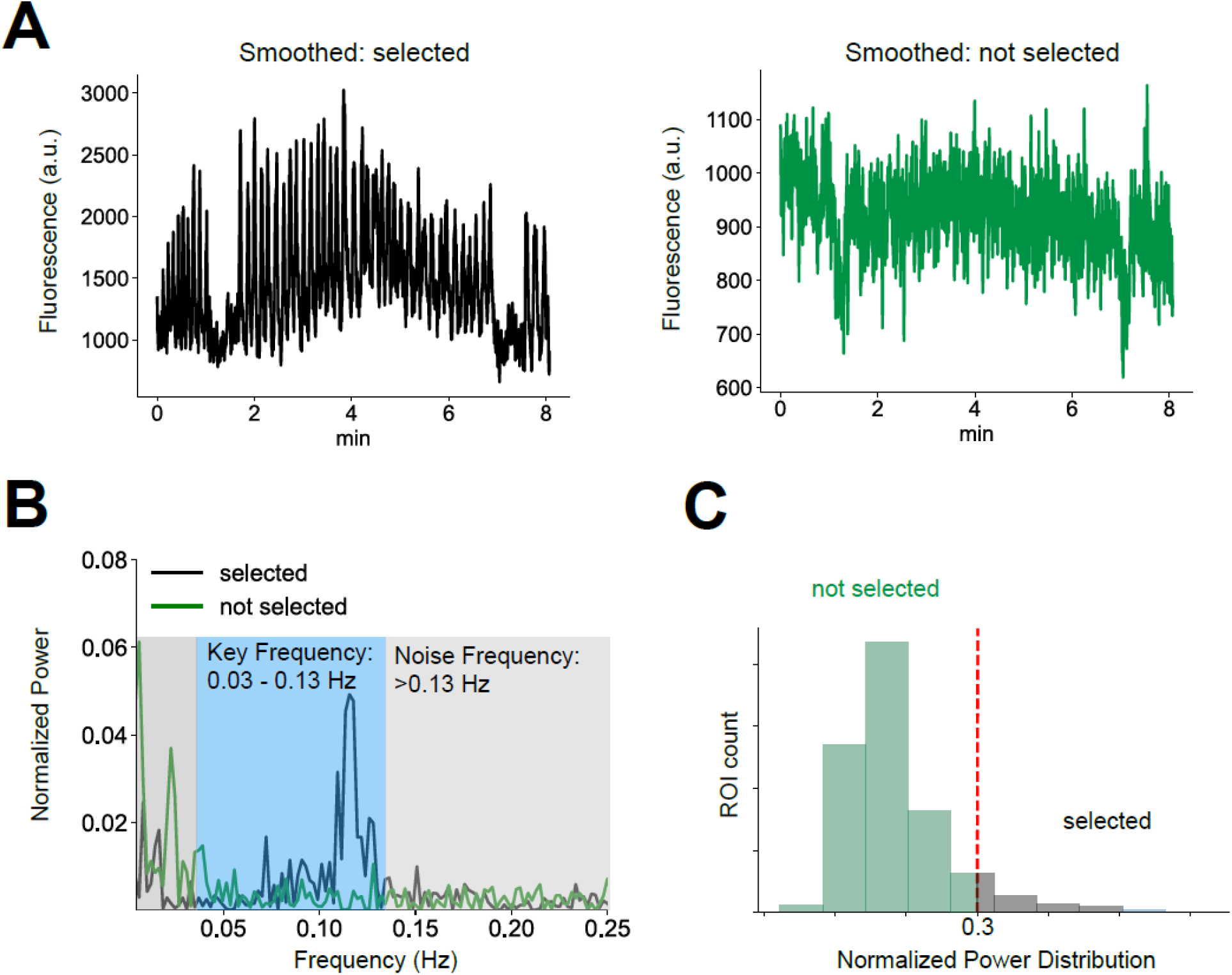
Frequency band-based axon ROI selection. **A**. Examples of selected (black) and discarded (green) ROIs in time domain after smoothing. **B**. Same ROIs in frequency domain. The x-axis displays the frequency of the FFT transformation, and the y-axis displays the normalized power. Note, the black ROI has a higher power in the frequency band of interest (shaded in blue) and lower power in the noise frequency (shaded in gray) while the discarded ROI is the opposite. **C**. The distribution of normalized band power within the frequency band of interest (0.03-0.13 Hz) for all example ROIs from a single FOV. The ROIs with lower power in the frequency band interest (below 0.3; dashed line) were discarded, and the ROIs with higher power in the band of interest (above 0.3) were selected.

### Time series anomaly detection for motion artifact identification

The previous step focuses on identifying transient-like patterns in our ROIs. However, the remaining ROIs might still contain traces that suffer from motion artifacts. X- and Y-axis movement are easily dealt with by motion correction algorithms like Suite2P. However, Z-axis movements may remain and are difficult to detect and remove. Typically, when a FOV moves along the Z-axis, many, if not all, ROIs are affected simultaneously. The effect of movement could be different in each ROI within the FOV (e.g. an ROI could drift in or out of the Z-plane). However, all affected ROIs would show synchronous abrupt changes (either an increase or decrease in signal) in fluorescence during a Z-axis shift. Thus, Z-axis shifts could be detected across the population of ROIs within a FOV by a sudden increase in coactivity strength and variance between ROIs. The goal here is to identify and remove periods of Z-axis shifts.

To achieve this, the first step is to z-score transform all ROI time-series traces to make them comparable. Real calcium transients should be stereotyped (as described above) and their occurrence should vary in time between ROIs, but z-axis shifts should cause synchronous signals that vary in terms of their shape (e.g. upwards or downwards). The distinct dynamics of real transients versus z-axis-induced signals, and the prediction that z-axis shifts will contribute to the largest variance in the moments shifting occurs compared to the rest of the time, should allow us to detect shifts using Principal Component Analysis (PCA) (**Fig. 4A**). We used the first principal component, which captures the largest variances in the data.

**Fig. 4.**
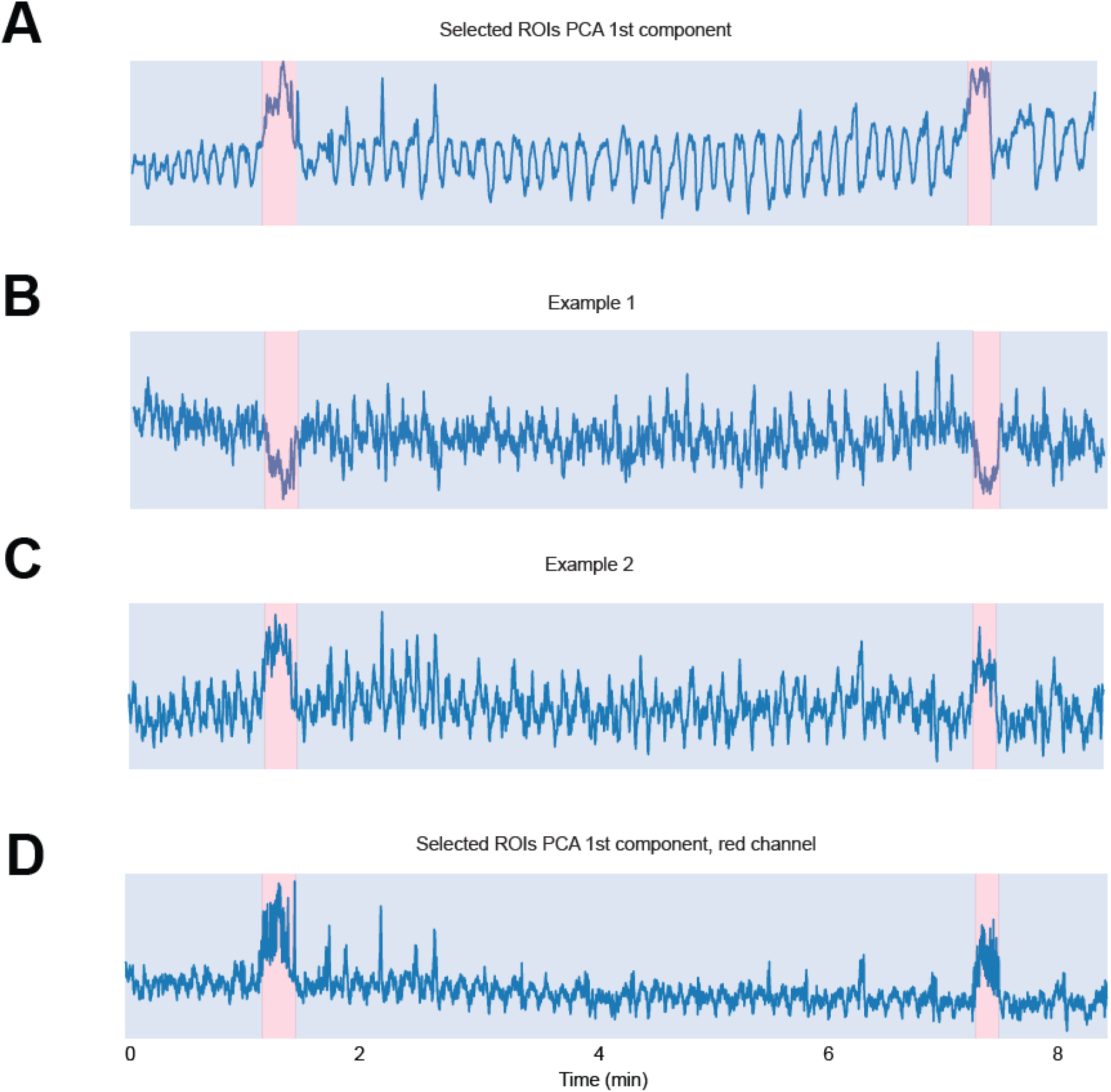
Identification of movement artifacts in time-series traces. **A**. First principal component on z-scored ROI activities. All selected ROI activities were first standardized to z-score and then dimension reduced to the first component of PCA. Bottom-Up algorithm was used to segment and detect anomalies (identified anomaly period shaded in red). **B & C**. Identified periods for potential movement artifacts were super-imposed on individual ROI activities for visual inspection. Two example selected ROI standardized activities are plotted and red shaded indicate potential periods of movement artifacts. All activities during the red shaded time period can then be ignored or removed from further data analysis. Note that the two example cells had activities in different directions but synchronous in time. **D**. Identified time periods for movement artifacts on red channel as further verification.

After applying PCA on z-scored ROI traces, we used a Bottom-up algorithm to segment and identify the time periods where a likely shift occurred[20]. Specifically, Bottom-up works by separating the time series data into a large number of very small segments of equal length and merging similar segments iteratively to bigger segments until met by a stopping criterion[21]. It is especially suitable for signals that contain complex and non-linear patterns with outliers and potentially multiple level shift changes, calculated by the algorithm. Using this method, we could detect shifts in the level of the baseline signal while filtering out noise and outliers. Each level shift change detected by the algorithm in the signal reflects a potential z movement shift artifact in the FOV (**Fig. 4A)**.

The bottom-up segmentation algorithm requires the number of change points it will detect to be input by the user. Using a larger number will lead to more periods detected by the algorithm and potentially more false positives and verse versa when using a smaller number. **Fig.4A** shows an example first PC which we visually inspected for potential Z-shifts. We used n=4 as the input for the bottom-up segmentation algorithm, which accordingly identified 2 periods of potential Z-shift. The user input at this stage requires consideration of neural dynamics of the imaged population. For instance, synchronous activity among large populations of ROIs can cause a level shift change in the first PC space. For regions of brain that have highly correlated activity, a static red fluorophore is needed to distinguish between movement artifact and synchronous activity. Our dataset was recorded from hippocampal CA3 axons during a spatial navigation task. While it is possible for groups of axons to be co-active, especially near to the rewards in the hippocampus[1, 2], the synchronous activity is unlikely to sustain for a long time (beyond a second or two). Thus, any prolonged level shift change in the PC space is highly likely to be driven by z-axis shift. Using a large number of n as input to the algorithm and then validating the results by measuring the duration of level-shift change reduces the possibility of having false positives.

In addition, a static red fluorophore could help to distinguish between population co-activity and z-axis shifts. The same z-score standardization and PCA can be applied to the red channel. **Fig. 4D** shows the PCA from the red channel from the same ROIs used in **Fig. 4A**. The red channel shows the same periods of elevated activity in the PC space identified in the green channel, confirming they are most likely caused by z-axis shifts. We can further validate the shifts using individual ROI activities (described below).

After finding periods of potential movement artifacts using PCA and Bottom-up algorithm, to visually confirm the validity of results, we underwent another manual visual inspection of the time-series traces from individual ROIs with labeled potential movement artifacts (**Fig. 4B and C)**. The two example ROIs illustrate that the movement artifact identified using the PCA method could be caused by z shifts from individual ROIs to be in the negative (**Fig. 4B**) or positive (**Fig. 4C**) direction, but the activities were synchronized during the labeled period, thus fitting our model of z-axis-induced signals. Multiple ROIs showing synchronized level shift in activity during the labeled period confirmed our findings of z-axis shift timing from PCA and Bottom-up algorithm. The frames during z-axis shifts can then be discarded across all ROIs within the FOV. Alternatively, if the FOV has shifted to a new Z-plane for a long time, our algorithm also allows for splitting the dataset into new sets of ROIs defined at the two planes to minimize data loss.

### Baseline correction and peak detection

To remove slow drifts in fluorescence caused by bleaching and scale the F signal to baseline, slow time scale changes in the fluorescence traces were removed by examining the distribution of fluorescence in a ∼20 second interval around each sample time point and subtracting the 8% percentile value. The resulting baseline corrected ΔF/F time-series traces were used to identify significant calcium transients. The initial steps in our pre-processing pipeline already detected ROIs with likely transients and removed motion artifacts. The following step was to further identify real transients from potential noise. To do this, peaks in the trace with a minimum amplitude of 0.12 ΔF/F, duration of 0.5 s, and prominence of 0.1 ΔF/F were identified. These regions we selected as “real” calcium transients and were included in the subsequent analysis.

### Clustering of the selected ROIs

Due to the density, coverage, and highly branched nature of axons and dendrites that can move in and out of the 2-photon imaging plane, multiple independent ROIs within a FOV could belong to the same cell. As a result, ROIs belonging to the same cell should have highly similar patterns of activity. Previously, one of the biggest challenges for axon clustering is a lack of ground truth labels to evaluate the performance of the clustering method. Failure to group axons or dendrites that branch from the same cell leads to overrepresentation of a single cell’s activity. Conversely, incorrect grouping of independent ROIs leads to loss of data and brings in artificially generated noise. In this section, we aim to compare two different clustering algorithms to combine axons and their corresponding performances when compared to ground truth labels.

Assuming that a high zero-phase cross-correlation between two ROIs reflects co-activation, we first performed a maximum correlation analysis across all ROIs over the entire time series (correlation matrix shape #ROIs x #ROIs). If the maximum correlation between a certain ROI with all the remaining ROIs exceeded a specific threshold (r = 0.8, see below for details), the ROI was likely to be part of a larger cluster of ROIs that belonged to the same axon. By contrast, if the maximum correlation between an ROI with all other ROIs was low, then the ROI was not likely to part of the same axon.

For the rest of the ROIs that could originate from the same axon, we calculated the pairwise correlation between ROIs and compared two different clustering methods, namely k-means and hierarchical clustering. To test whether the clustering methods were doing a good job, we compared their ROI clustering results to “ground truth data”, i.e. ROI groupings that were highly likely from the same axon. To obtain ground truth data in our experimental setup, animals ran through two different virtual reality environments while imaging hippocampal CA3 axons in the CA1. A feature of hippocampal cellular activity is their spatial firing and their ability to remap their spatial firing fields upon exposure to a novel environment, meaning that while it is possible for distinct place cells to have similar spatial firing within one environment, it is highly unlikely for them to also fire in a similar pattern in more than one environment[22]. Thus, if the pairwise correlation between ROIs was consistently above 0.7 across two distinct environments, we considered those pairs to belong in the same axon group. This served as our “ground truth data” in which our clustering methods could be compared.

We evaluated the similarity between the ground truth data and our assigned clusters using two different metrics: the Adjusted Mutual Information (AMI) score and the Silhouette score. While the AMI score measured the similarity between the true cluster labels (i.e., > 0.7 correlations between two ROIs in both of our virtual environments) and the labels assigned by the clustering algorithm (e.g., k-means or hierarchical clustering), adjusting for chance, the Silhouette score evaluated the quality of the clusters themselves without the need for ground truth labels. The AMI score ranged from 0 to 1, where a score of 1 indicated perfect agreement between true and predicted cluster labels. The Silhouette score ranged from −1 to 1, with higher values indicating better clustering results. This metric assessed the separation of clusters based solely on the clustering output produced by clustering algorithms. Specifically, a high Silhouette score indicates clear separation between cluster groups, that objects within a cluster were close to each other and far away from objects in other clusters. A high performance in either metric suggests that the clustering results were reliable, and the number of clusters that maximize these metrics represents the best-fit grouping assignments for the remaining ROIs. Because not every axon dataset will have ground truth labels to calculate AMI, we compared the consistency between AMI and Silhouette scores to test if Silhouette score is sufficient to lead to a similar clustering result without the need for ground truth data.

To estimate the number of clusters, K number of clusters was examined from 2 to 50% the of the selected ROIs. For both example mice, Silhouette scores and AMI scores converged (contained a maximum number that represent best performance) and reached optimal K for clustering (see **Fig. 6A**, left panel). In addition, both performance metrics (AMI and Silhouette) returned a similar number K, which suggests that hierarchical clustering was efficient with or without true labels of clusters.

**Fig. 5.**
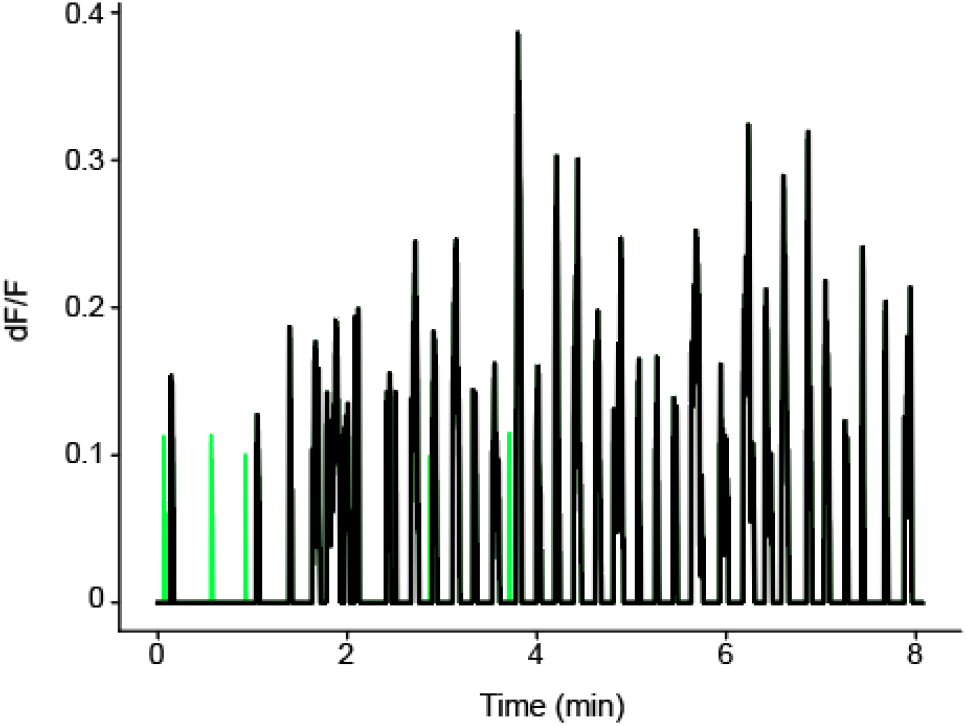
Baseline correction and peak detection. All ROI activities were scaled to baseline to create ΔF/F traces. Baseline corrected ΔF/F traces across time were generated using sliding window of ∼20 seconds and the 8% percentile value within the sliding window was subtracted from each timepoint. Across the baseline corrected ΔF/F traces, peaks were calculated using minimum height, distance, and prominence values. Of all potential peaks, only detected peaks (black) were kept and all other activities (green) were forced to 0.

**Fig. 6.**
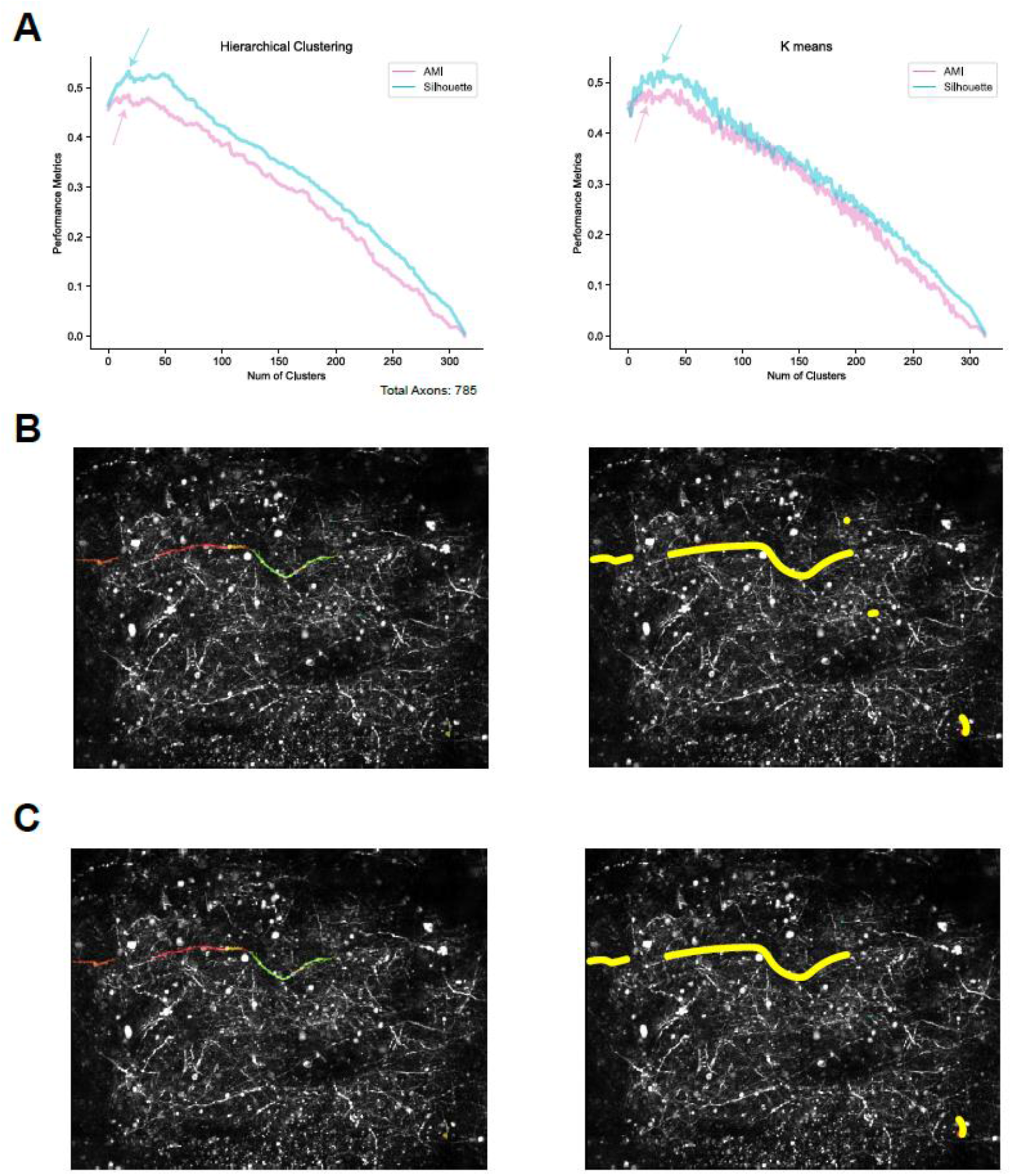
Comparison of clustering methods used for grouping ROIs. **A.** Clustering performance metrics with hierarchical clustering and K-means clustering. The left panel shows that hierarchical clustering converged to a similar optimal number of clusters (peak pointed by the arrows) for both performance metrics, Silhouette scores and AMI. The right panel showed that K-means clustering converged to a similar optimal number of clusters for both performance metrics, Silhouette scores and AMI. **B.** Example grouping from hierarchical clustering transposed to original FOV. Left: each color represents an individual ROI in the group. Right: all ROIs in the same group highlighted in yellow **C.** Same grouping from “ground truth” data.

In addition to hierarchical clustering, the k-means algorithm can also be used to identify clusters of high correlations between pairs of ROIs. Like hierarchical clustering, k-means uses the full correlation matrix between all pairs of ROIs as input (citation needed here). Our main goal was to identify the optimal number of clusters, k, that best explains the covariance structure of the ROIs. We applied the k-means algorithm with a predefined number of clusters, ranging from 2 to half the number of ROIs (n). The algorithm partitions the samples into k clusters based on similarity, iteratively assigning each sample to the nearest cluster centroid and updating centroids to minimize the within-cluster sum of squares. This process continues until convergence, resulting in clusters where correlations between ROIs are maximized within each cluster and minimized between clusters. Notably, the optimal number of clusters identified by k-means closely matched the results from hierarchical clustering. Therefore, we conclude that both methods reliably estimate optimal clustering.

### Summary of the SUBPREP

A sample paradigm of our SUBPREP pipeline is displayed in **Fig. 7**, with all the significant steps and an example ROI on the right side of the flowchart. The first step, pre-processing, was applied to all ROIs and the goal was to smooth the data across time. After pre-processing, the ROIs with higher power in the key frequency band for transients were selected with band-pass filtering methods (0.03 to 0.13 Hz). Then, to identify movement artifacts, we standardized and used PCA to reduce the dimensions in the population activity. We used a Bottom-up algorithm to identify anomalies in signal of the reduced PC space. We confirmed the results by imposing the labeled movement artifact periods onto individual ROIs and visually inspected their activity. Confirmed periods of movement artifact were removed from all ROIs in the FOV. The remaining ROI signals, highly likely to contain transients from stable FOVs, were identified by thresholding signal prominence which allowed detection of transient peaks. Lastly, to identify the potential clustering of the ROIs belonging to the same axon, we performed two methods, hierarchical clustering and k-means clustering. Both methods were effective in identifying the optimal number of clusters in the pairwise ROI correlation matrices and resulted in similar grouping compared to the ground truth labels.

**Fig. 7.**
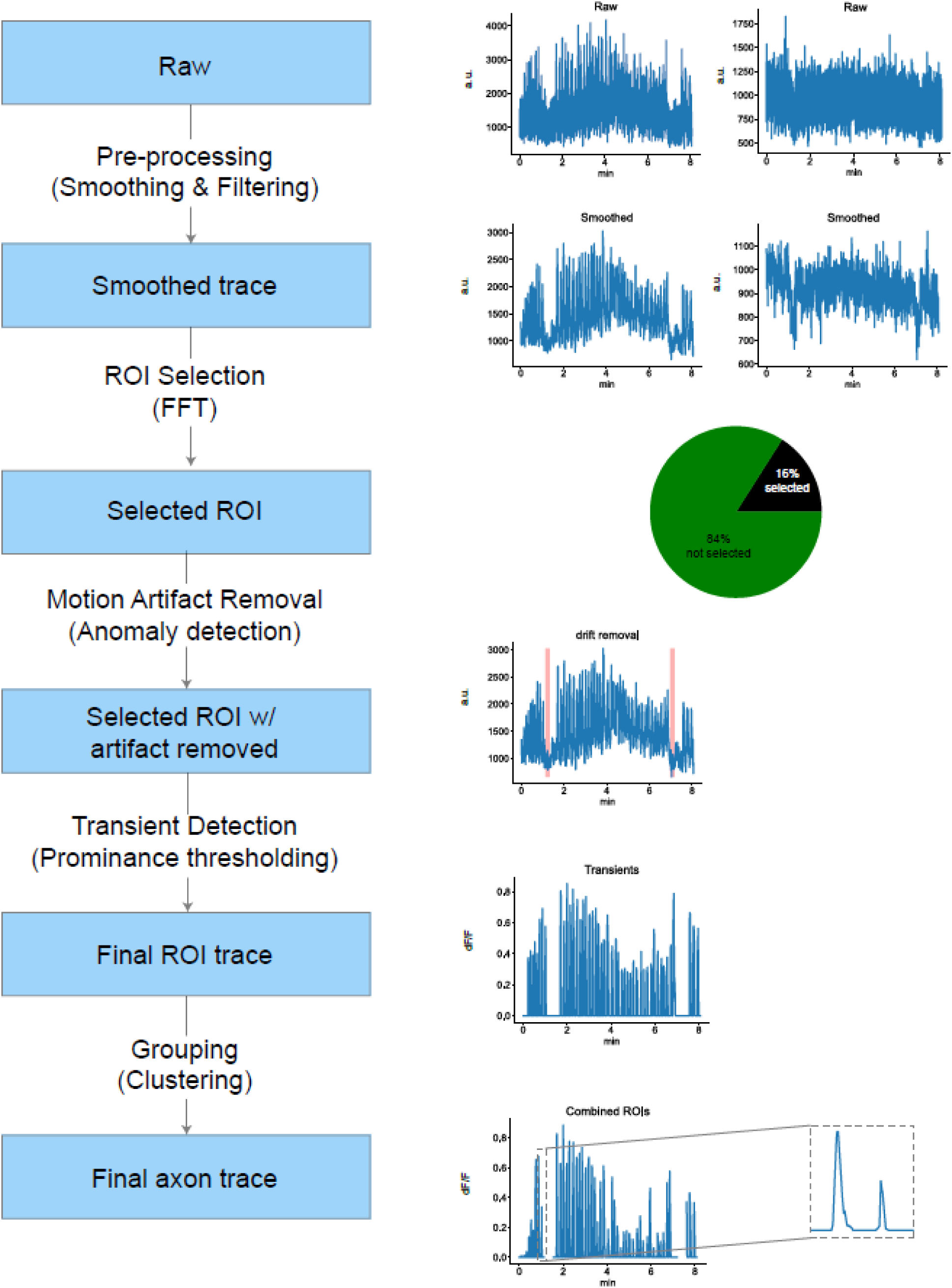
SUBPREP pipeline summary.

## DISCUSSION

In this study, we developed and validated a computationally efficient preprocessing pipeline for subcellular signal detection, movement artifact identification, and ROI grouping in two-photon calcium imaging data. Our approach addresses several key challenges associated with analyzing subcellular structures, such as low signal-to-noise ratio (SNR), movement artifacts, and difficulty in grouping subcellular ROIs from the same neuron. Through the application of frequency-based filtering, Principal Component Analysis (PCA) and level change detection for motion artifact identification, and clustering algorithms for ROI grouping, our pipeline provides a robust framework for extracting physiological signals from subcellular structures during behavior.

Suite2P and SIMA, designed for detecting somatic activity, provide robust frameworks for initial subcellular data processing[16, 23], but additional steps are required as subcellular datasets are inherently noisy and sensitive to Z-plane movements because of their small size relative to somas. Our processing pipeline offers a systematic approach to sub-select ROIs from the initial pool identified by Suite2P by leveraging the unique frequency domains of calcium transient dynamics. By employing Fast Fourier Transform (FFT) and band-pass filtering, we were able to isolate ROIs likely to contain true calcium transients, allowing accurate signal detection in noisy subcellular datasets. Other methods have been used for this step, such as manual ROI selection and convolutional neural network (CNN)-based approaches, but these are either labor-intensive and prone to bias, or require extensive ground truth data for training[1, 2, 12]. A second, “static”, fluorescence channel has often been used to subtract noise from an “active” channel, usually from GCaMP signals, but this can introduce artifacts as noise is not perfectly mirrored across channels. For instance PMTs are a source of noise and separate PMTs are used for 2 channel imaging[19], and the fluorophores themselves are distinct sources of noise. Our approach offers an unbiased and computationally cheap method for subcellular ROI identification that does not require dual-color imaging, thereby simplifying experimental design and reducing potential artifacts from channel subtraction.

Accurately identifying periods of Z-plane shifts is always important but is amplified when imaging subcellular ROIs versus somas because axons and dendrites are relatively small and have elaborate structures that can be found in and out of the imaging plane in complex branching patterns. A small movement in Z can result in a distinct set of subcellular ROIs being imaged. After identifying ROIs with potential transients, our pipeline effectively identifies Z-plane movement artifacts by utilizing PCA and a Bottom-Up Segmentation algorithm to detect artifacts based on the first principal component. This allows for the analysis of data collected from a stable imaging plane by excluding time points collected from different planes. Additionally, if a prolonged Z-shift to a new imaging plane occurs, the new Z-plane can be identified and used for data analysis of a distinct set of subcellular ROIs during this period. This limits data loss from technically challenging subcellular imaging experiments.

Despite the strengths of our approach, several limitations remain. For instance, our method’s reliance on frequency-based filtering requires careful consideration of the smoothing filters applied prior to FFT, as these can impact the key frequency band of real transients and thus the selection of ROIs. Additionally, while our PCA-based motion artifact identification effectively detects Z-plane shifts, it still requires user input to define the number of change points, which introduces potential bias. Automating this process or developing more sophisticated algorithms to reduce user dependency could further improve the robustness of the pipeline.

Future work should integrate our SUBPREP toolbox into existing preprocessing calcium imaging software, specifically Suite2P. This would provide multiple benefits for the user. One benefit would be that the user interface and real-time visual displays in Suite2P could be utilized for SUBPREP to show the user which ROIs in the FOV contain real transients or which segments of the time-series movies have potential motion artifacts. It would also streamline analysis allowing one software package that can be used for cellular and subcellular preprocessing. Other future work should focus on expanding the validation of our approach across different brain regions and experimental conditions will be crucial for generalizing its applicability.

SUBPREP offers a standardized approach for analyzing axonal and dendritic activity by simplifying the process of ROI selection, artifact removal, and ROI grouping, providing a practical and effective solution to several key challenges in the field of sub-cellular imaging. We hope our publicly available processing pipeline, SUBPREP, is used by the field to help standardize the extraction of physiological signals from subcellular structures, enabling more accurate and reproducible studies of subcellular activity across labs. This would lead to more robust insights into the mechanisms of information transmission and integration in the brain, ultimately advancing our understanding of neural circuitry and its role in behavior.

## ACKNOWLEDGEMENTS

We thank Seetha Krishnan, PhD for feedback on the manuscript and all members of the Sheffield lab for valuable discussions that helped shape the manuscript.

## AUTHOR CONTRIBUTIONS

A.J. and M.S. conceived and designed the experiments. A.J. collected the data. A.J. and C.Z analyzed the data and wrote the SUBPREP software. A.J., M.S., and C.Z. interpreted the data and wrote the manuscript. M.S. supervised the research and obtained funding.

## DATA AVAILABILITY

Raw imaging data is large and not feasible for upload to an online repository but is available upon request to the lead contact at. Processed source data for all figures and associated statistical analysis are provided will be provided in the final version of the paper.

## CODE AVAILABILITY

SUBPREP software used for subcellular data analysis is available here: (https://github.com/anqijiang/subprep).

